# Dermaseptin DMS-DA6, a promising alternative to conventional antibiotic in the treatment of *N. brasiliensis* induced actinomycetoma

**DOI:** 10.1101/2025.06.16.659840

**Authors:** Duarte-Mata Diana Ivonne, Vázquez-Marmolejo Anna Velia, Alemán-Navarro Estefania, Piesse Christophe, Rosenstein Yvonne, Auvynet Constance, Salinas-Carmona Mario Cesar

**Author notes:** **Correspondance:** Dr. Mario Cesar Salinas-Carmona: Av. Gonzalitos 235, Mitras Centro, 64460 Monterrey, N.L., Dr. Constance Auvynet: Av. Universidad 2001, Chamilpa, 62210 Cuernavaca, Mor.

## Abstract

Actinomycetoma is a chronic infectious disease recognized by the World Health Organization as a neglected tropical disease. In the Americas, the most common etiologic agent is the Gram-positive bacterium, *Nocardia brasiliensis*. It is a facultative intracellular pathogen that can multiply and survive within macrophages, evading microbicidal mechanisms by inducing an immunosuppressive environment. Current antibiotic treatments are expensive, prolonged, and toxic, and bacterial resistance has been reported. Host defense peptides, known for their bactericidal and immunomodulatory effects, and their ability to induce poor bacterial resistance due to their direct effect on the bacterial membrane, represent a novel therapeutic approach. Recently, DMS-DA6, a dermaseptin isolated from the Mexican tree frog, *Pachymedusa dacnicolor*, has been shown to exert strong activity against Gram-positive bacteria, including multidrug-resistant strains. Our study evaluated the effects of DMS-DA6 in a *N. brasiliensis*-induced actinomycetoma mouse model in comparison with the ones observed with the conventional antibiotic linezolid. Infected mice were treated either twice a week with this peptide at a dose of 12.5mg/kg or every 12 hours with linezolid at a dose of 25 mg/kg over a three-week period. Our findings suggest that treatment with DMS-DA6 is more effective in resolving the disease than linezolid, as it induces a similar reduction in the volume of inflammation and bacterial load in the infected area at a lower dose and with fewer injections. These findings highlight the potential of DMS-DA6 as an innovative addition to current therapy.

**Author Summary:** Actinomycetoma is a chronic infectious disease induced by Gram-positive bacteria and listed as a neglected tropical disease that produces severe tissue deformation, and family, social, and economic problems. Current treatment takes years, is expensive, and induces undesirable effects and bacterial resistance, leading to amputation of extremities in many cases. There is thus an urgent need to develop new therapeutics, and host defense peptides could be promising candidates. In the *Nocardia brasiliensis* actinomycetoma mouse model, we demonstrated that the host defense peptide DMS-DA6 was more effective than the classical antibiotic linezolid in decreasing the bacterial load and inflammation in the infected footpad, paving the way for its incorporation in human trials. We thus propose the use of DMS-DA6 as an alternative therapy before amputation of the infected extremity.

## Introduction

Actinomycetoma is an infectious chronic subcutaneous granulomatous disease with a severe inflammatory component, that is slow and progressive in its development, and affects tissues by deforming and destroying them (1). The causative agents are facultative intracellular bacteria of the Actinomycetales order. In Mexico, actinomycetoma is responsible for around 97% of the infections, with *Nocardia brasiliensis* being the most prevalent causative agent (65%) (2).

Even if the pathogenic mechanism remains unclear, it has been shown that in the chronic phase of the infection, an immunosuppressive environment with high IL-10 and low IL-6/IFNγ levels promotes inhibition of phagocytosis and apoptosis of macrophages and differentiation of macrophages and dendritic cells into foamy cells which favors the survival and multiplication of the pathogen into these foamy cells (3,4).

The standard treatment for actinomycetoma is trimethoprim/sulfamethoxazole (TMP/SMX) (5). If resistance or side effects like ototoxicity or nephrotoxicity arise, the treatment is stopped, and linezolid is recommended, as it is effective against all Nocardia species (6,7). However, current treatments, including linezolid, are often lengthy, exhibit severe side effects, and have high relapse rates (8). Consequently, the development of safer and more effective therapies is of the greatest importance.

Host defense peptides (HDPs) are potential candidates as they play a crucial role in controlling infections through direct bactericidal action against a wide range of bacteria, including multi-resistant strains, and through modulation of the immune system (9). However, due to their unique structures and direct action on the microbial membrane, they have been shown to be less prone to inducing bacterial resistance than conventional antibiotics (10). Among all peptide-producing organisms, HDPs isolated from frogs have been extensively identified and characterized (11). Among them, Dermaseptins (DRS) are a family of α-helical-shaped polycationic peptides found in the skin secretions of Latin American Hylid frogs (12). The dermaseptin DMS-DA6 isolated from the skin of the Mexican frog *Pachymedusa dacnicolor* exhibits remarkable bactericidal activity against Gram-positive bacteria, including multi-resistant strains, with no hemolytic activity (13).

In this study, the effects of DMS-D6 on disease progression and bacterial load were evaluated in a mouse model of actinomycetoma induced by *N. brasiliensis* and compared to those observed after treatment with linezolid. Our results demonstrate the potential benefits of using the HDP DMS-DA6, as it controls disease progression by reducing the bacterial load at a lower dose and with fewer injections than linezolid while also modulating the immune system, leading to a diminution of the volume of the infected footpad. Our findings pave the way for the future use of these molecules in human cases.

## Materials and Methods

### DMS-DA6 Peptide

The peptide was synthesized by automated solid-phase peptide synthesis using Fmoc/tBu (9-Fluorenylmethoxycarbonyl/tertio-butyl) chemistry on an ABI 433A peptide synthesizer (Applied Biosystems) and purified by RP-HPLC (C18 Xbridge Waters, 5 μm 19 × 50 mm) using a solvent system composed of water containing 0.1% trifluoroacetic acid (TFA) as solvent A and acetonitrile containing 0.1% TFA as solvent B. The column was eluted at 10 mL. min-1 with a 35–60% linear gradient of solvent B for 10 min. Peptide purity was assayed by analytical HPLC (C8 Ace, 300Å, 5 μm, 250 × 4.6 mm, Ait France). Identities of the peptides were confirmed by MALDI-TOF mass spectrometry (MS Voyager Applied Biosystems). Considering the presence of one tryptophan residue in the sequence, concentrations were determined by UV spectroscopy (Nanodrop, Fischer Scientific), assuming an extinction coefficient for Tryptophan ε280 = 5 600M-1.cm-1.

### Mice

Ten- to twelve-week-old female BALB/c mice were used for the present study. Animals were maintained under BSL2 conditions at the Service of Immunology facility for rodents. Mice received sterile water and Purina rodent food *ad libitum*. Protocols for animal care and infections were developed according to the International Review Board regulations and Mexican regulations (NOM-062-ZOO- 1999) and approved by the Bioethics Committee of the Facultad de Medicina at the Universidad Autónoma de Nuevo León (Registration number: IN23-00007).

### DMS-DA6 and linezolid treatment for actinomycetoma by *N. brasiliensis* in BALB/c mice

Actinomycetoma infection was induced in mice by injecting 10^6^ CFU of *N. brasiliensis* ATCC 700358 into a rear footpad as previously described (14). A total of 20 mice were used in this study (5 mice per treatment). The first dose of DMS- DA6 or Linezolid was administered in the chronic phase of infection. DMS-DA6 peptide was injected intramuscularly at a concentration of 12.5 mg/kg body weight twice a week for three weeks. Linezolid was injected subcutaneously at a concentration of 25 mg/kg body weight twice daily for four weeks. Uninfected and infected mice used as controls received 20 µL of saline solution by intramuscular injection.

Survival and animal weight were recorded individually every other day. The volume of inflammation in the infected footpad was recorded with the use of a vernier caliper, using the ellipsoid equation (width x height x length).

Peripheral venous blood was collected for flow cytometry analysis, and serum samples were taken for cytokine quantification after three or four weeks of treatment. Controls and treated animals were euthanized (n=5) to prepare cell suspensions of infected footpads and spleen for cytometry. Footpads were homogenized to determine bacterial load and digested with collagenase XI (0.7 mg/mL, Sigma) in RPMI for 30 minutes at 37°C. The cell suspension was filtered with a 70-µm cell strainer (BD Biosciences) and treated with a lysing solution before staining for cytometric analysis.

### Flow cytometric analysis

Cell suspensions from the footpad, spleen and blood containing 10^6^ cells/ml from each mouse were stained with the following monoclonal antibodies (mAbs): anti- CD3 (APC-Cy7), anti-CD4 (Pacific blue), anti-CD8 (V500), anti-CD11b (Alexa fluor 647), anti-Ly6G (PE-Cy7) and anti-F4/80 (Alexa fluor 488). All mAbs were from BD Biosciences. Cell suspensions were incubated with mAbs at 4°C for 30 min in the dark, washed, and suspended in 500 µL of PBS. Data acquisition and analysis were completed using a Flow Cytometer LSR Fortessa (BD Biosciences) and DIVA software v8.0 (BD Biosciences, San Jose, CA), respectively. The acquisition of at least 100,000 total events in the cell gate was used to report percentages.

### Quantification of plasma cytokines

Cytokines were measured in the sera from control or infected and treated mice with the LEGENDplex Mouse Inflammation Panel 13-Plex system (BioLegend), following the manufacturer’s instructions for the quantification of the cytokines and chemokines known to be implicated in the chronic phase of the infection: IL-1, IL-6, IFNγ, TNF-α, IL-10 and MCP-1 by flow cytometry. Sample acquisition was performed with a BD FACSCanto II flow cytometer (BD Biosciences) and the BD FACSDiva software v9.0.1. Data were analyzed with the LEGENDplex Data Analysis Software from BioLegend (https://legendplex.qognit.com).

### *N. brasiliensis* bacterial load

Serial dilutions of footpad homogenized footpad tissues were used to determine bacterial loads by plating on brain heart infusion agar and incubating for one week at 37°C. Colony-forming units are expressed per milliliter of suspension.

### Statistical analysis

Statistical analyses were performed using GraphPad Prism software v.8. Differences between groups were evaluated by one-way ANOVA followed by Tukey’s post-test, Mann-Whitney test, and Unpaired t test. *P* values of <0.05 were considered significant.

## Results

### DMS-DA6 reduces the volume of the infected footpad in *N. brasiliensis* actinomycetoma

A chronological overview of the administration of the antimicrobial peptide and antibiotic to mice infected with *Nocardia brasiliensis* is presented in Fig. 1A. After administering DMS-DA6 or linezolid for three weeks during the chronic stage of infection, 100% survival was noted (Fig. 1B). A 5% variation in body weight was observed in the infected untreated group as compared to the non-infected group. This variation tended to decrease after treatment with DMS-DA6 or linezolid, although the decrease was not statistically significant (Fig. 1C). Furthermore, the administration of DMS-DA6 did not cause any irritation or clinical signs indicative of acute toxicity (data not shown).

**Figure 1.**
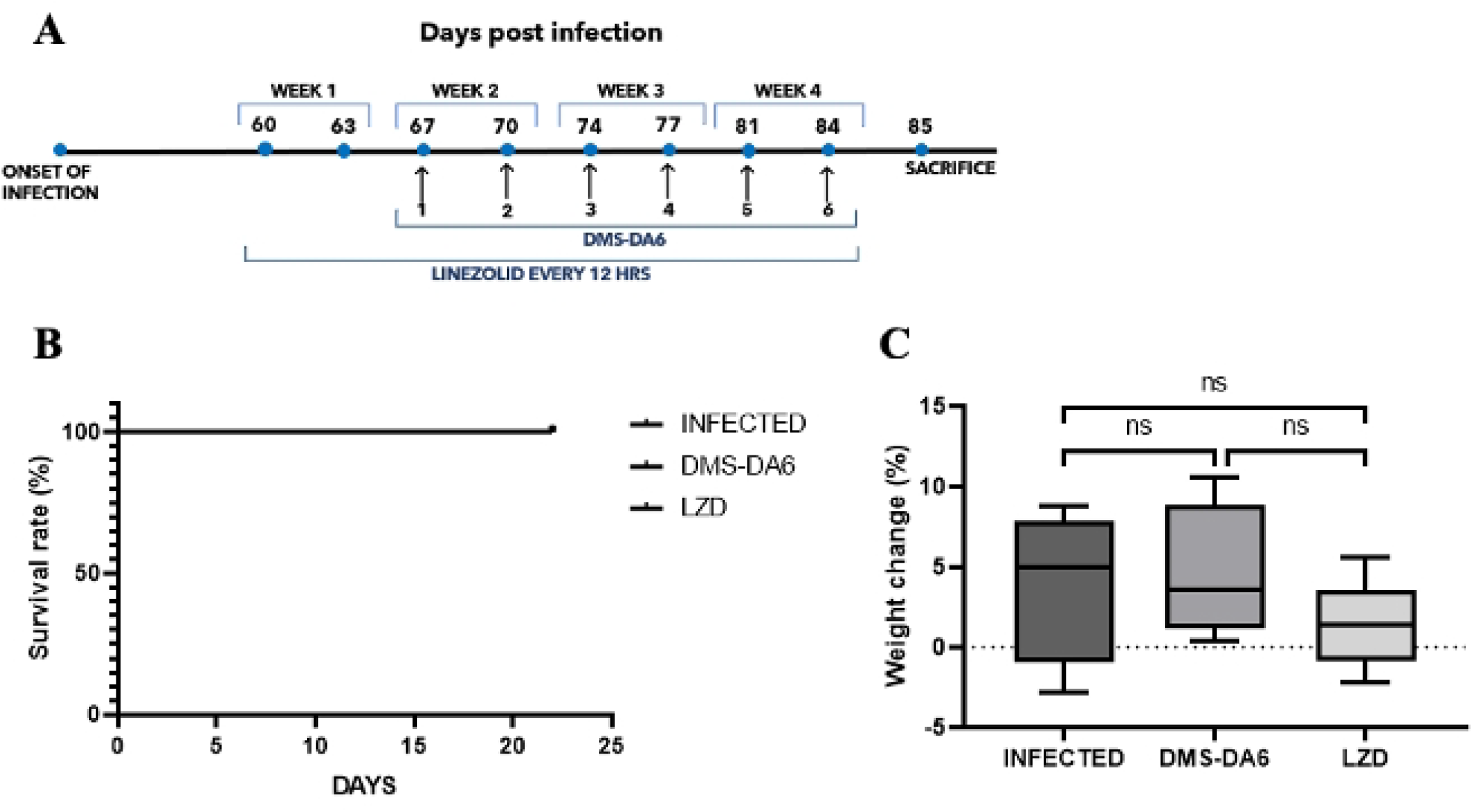
*In vivo* administration of DMS-DA6 does not affect the survival or weight of mice with actinomycetoma. (A) Treatment scheme of the administration of the DMS-DA6 (12.5mg/kg) and Linezolid (25mg/kg) in mice at the chronic stage of infection by *N.brasiliensis*. (B) Survival rate after treatment with DMS-DA6 and linezolid (n=5). (C) Changes in body weight were assessed during the treatment. Unpaired t test (n=5) ns: p>0.05.

The inflammation of the infected footpad from untreated mice and those treated with DMS-DA6 or linezolid is shown in Fig. 2. As expected, the volume of the footpad in the untreated infected group increased by up to 50% over the weeks, while treatment with DMS-DA6, at a dose of 12.5 mg/kg twice a week, or that of Linezolid, at a dose of 25 mg/kg every 12 hours, gradually reduced the volume of the footpad, starting from the second week onward (Fig 2B). Notably, even though the reduction in volume of the infected footpad induced by DMS-DA6 is similar to that resulting from the linezolid treatment, DMS-DA6 was administered at a lower dose and with fewer injections, suggesting a greater efficacy of DMS-DA6.

**Figure 2.**
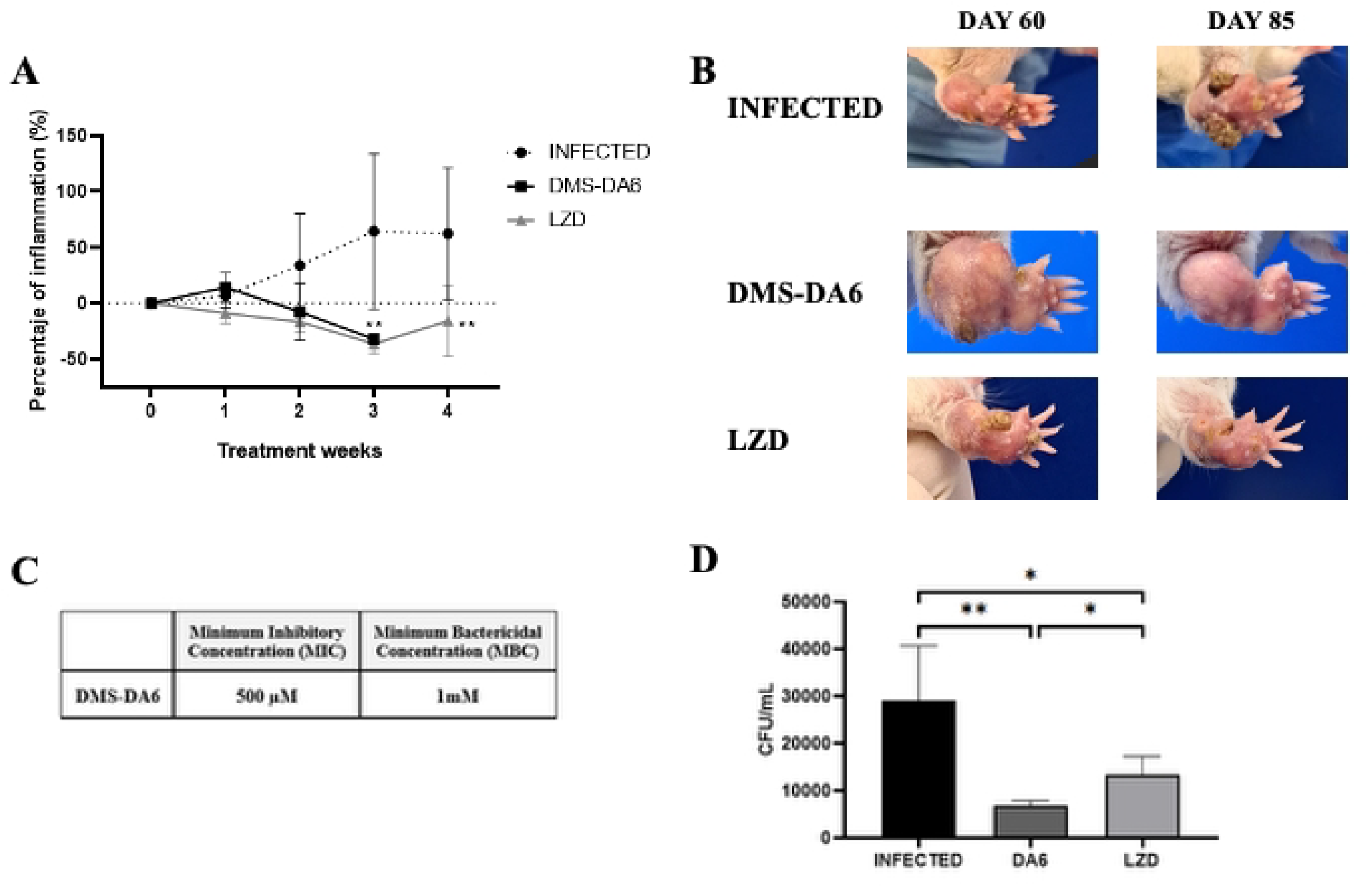
*In vivo* administration of DMS-DA6 reduces the volume and the bacterial load in the infected footpad. **(A)** Volume changes of the infected footpad in non-treated and linezolid or DMS-DA6 treated mice during were measured during the treatment. (B) Clinical course of the infected footpad is shown before and after treatment with DMS-DA6 or Linezolid. (C) Minimum inhibitory concentration (MIC) and minimum bactericidal concentration (MBC) of DMS-DA6 against *N. brasiliensis*. (D) Colony forming units (CFU) were determined in the infected footpad of linezolid or DMS-DA6 treated mice. (n=5) 2Way ANOVA, Tukey’s multiple comparisons. Unpaired t test *p<0.05 **p<0.01.

### DMS-DA6 exhibits stronger antibacterial activity in comparison to linezolid

The antibacterial activity against *Nocardia brasiliensis* was assessed *in vitro*. As shown in Figure 2C, DMS-DA6 inhibited bacterial growth with a MIC of 500 µM and induced bacterial death with an MBC of 1 mM. The bacterial load in the footpad was then analyzed in both the treated and untreated infected groups. As shown in Figure 2D, DMS-DA6 and linezolid significantly decreased the bacterial load in the infected footpad compared to the untreated group. Interestingly, even if the MIC and MBC observed for DMS-DA6 *in vitro* were relatively high, DMS-DA6 exhibited significantly stronger antibacterial activity compared to linezolid, with fewer CFU in mice treated with the peptide than with the antibiotic.

### DMS-DA6 modulates IL-10, IL-1*α*, and IL-6 cytokine levels at the chronic stage of actinomycetoma infection

In the chronic stage of infection, an increased production of IL-10 and a decreased production of IFNγ, IL-6, IL-1, and TNF-α play an important role in sustaining chronic infection by enabling the survival of the pathogen within foamy macrophages. We thus evaluated whether DMS-DA6 could modulate the production of these cytokines. Interestingly, we observed that the treatment with DMS-DA6 significantly decreases the serum levels of IL-10 (Fig. 3A) and IFNγ (Fig. 3B) while increasing those of IL-1α (Fig. 3C) and IL-6 (Fig. 3D) compared to the untreated infected group. Although not significant, a decrease in TNFα level (Fig. 3E) was observed. In contrast, linezolid administration did not result in any alterations in the serum levels of these cytokines, except for IFN-γ.

**Figure 3.**
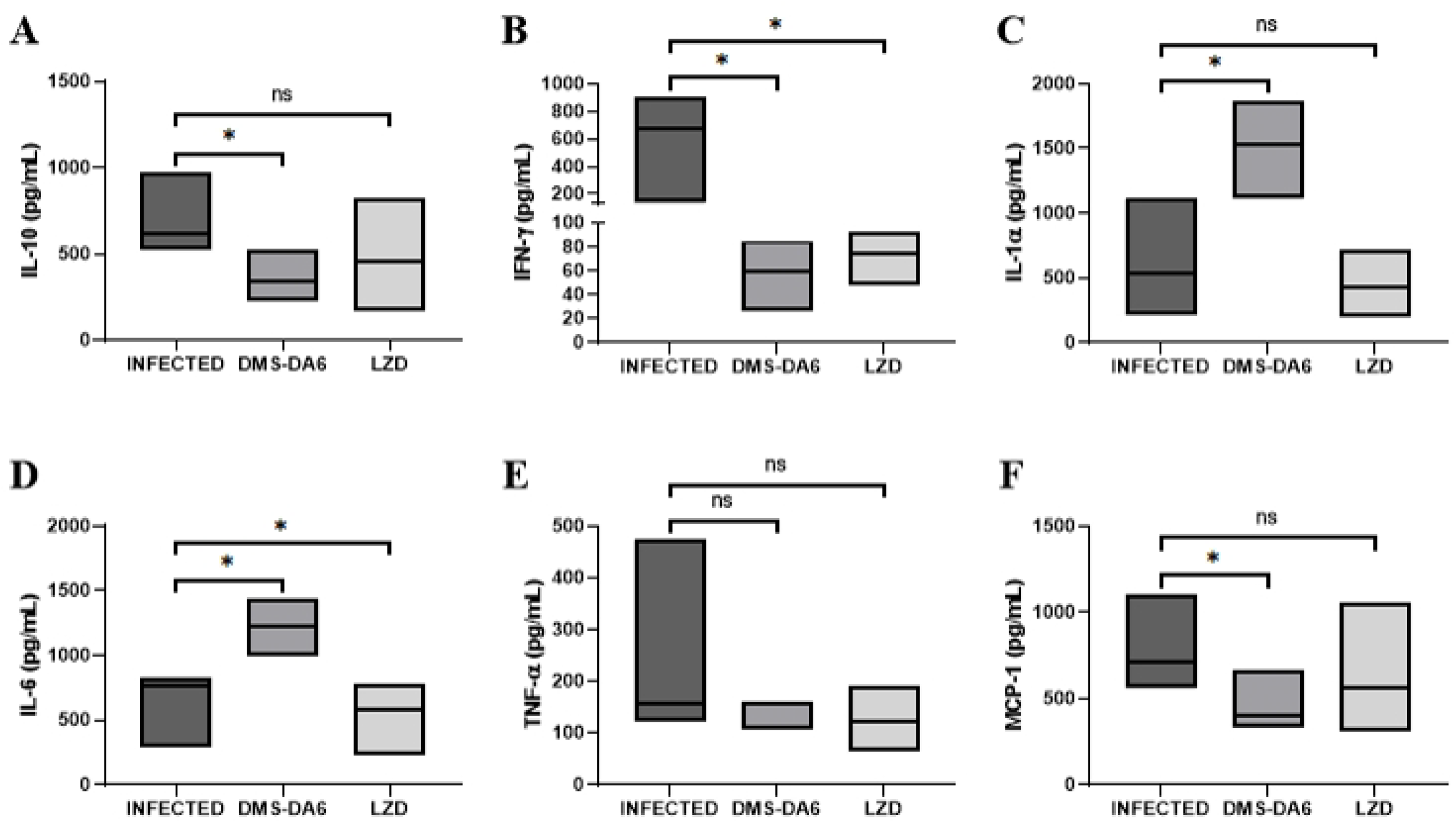
DMS-DA6 modulates IL-10, IL-1*α*, and IL-6 cytokine levels at the chronic stage of actinomycetoma infection. Levels of IL-10 (A), IFN-γ (B), IL-1α (C), IL-6 (D), TNF-α (E) and MCP-1 (F) in the serum of actinomycetoma untreated or DMS-DA6 and linezolid treated mice were evaluated by flow cytometry and analyzed with DIVA Software. (n=5) Mann-Whitney test and Unpaired t test *p<0.05 ns: p>0.05.

As macrophages are important for Nocardia survival, we also assessed the effect of DMS-DA6 on MCP-1 level, a chemokine known to promote the migration of classical monocytes that differentiate into macrophages in inflamed tissue. We observed a significant decrease in this chemokine level after treatment with DMS-DA6, consistent with the reduction of bacterial load and inflammation in the footpad (Fig. 3F). No effect on MCP-1 level was observed after treatment with linezolid.

Collectively, these results suggest that DMS-DA6 modulates the production of cytokines implicated in the survival of Nocardia and thus potentiates the direct antibacterial effect induced by this peptide.

### At the chronic stage of infection, DMS-DA6 treatment boosts the percentage of CD4+ and CD8+ T lymphocytes in the infected footpad

Since CD4+ and CD8+ T lymphocytes have been shown to contribute to controlling Nocardia infection, we assessed how DMS-DA6 modulates these populations. Indeed, it has been demonstrated that the proportion of CD4+ and CD8+ lymphocytes decreases in the chronic phase of intracellular infections (ref). Interestingly, the peptide significantly increased the percentage of CD4+ T lymphocytes (Fig. 4A) and tended to increase that of CD8+ T lymphocytes (Fig. 4B) in the infected footpad. These results align with observations regarding IL-10 production, where a decrease in IL-10 production correlates with an increase in the percentage of CD4 and CD8+ T lymphocytes (15). No effect was observed on CD4+ and CD8+ T lymphocytes after treatment with linezolid. Macrophages and neutrophils were not affected by any of the treatments (Fig. 4C and D).

**Figure 4.**
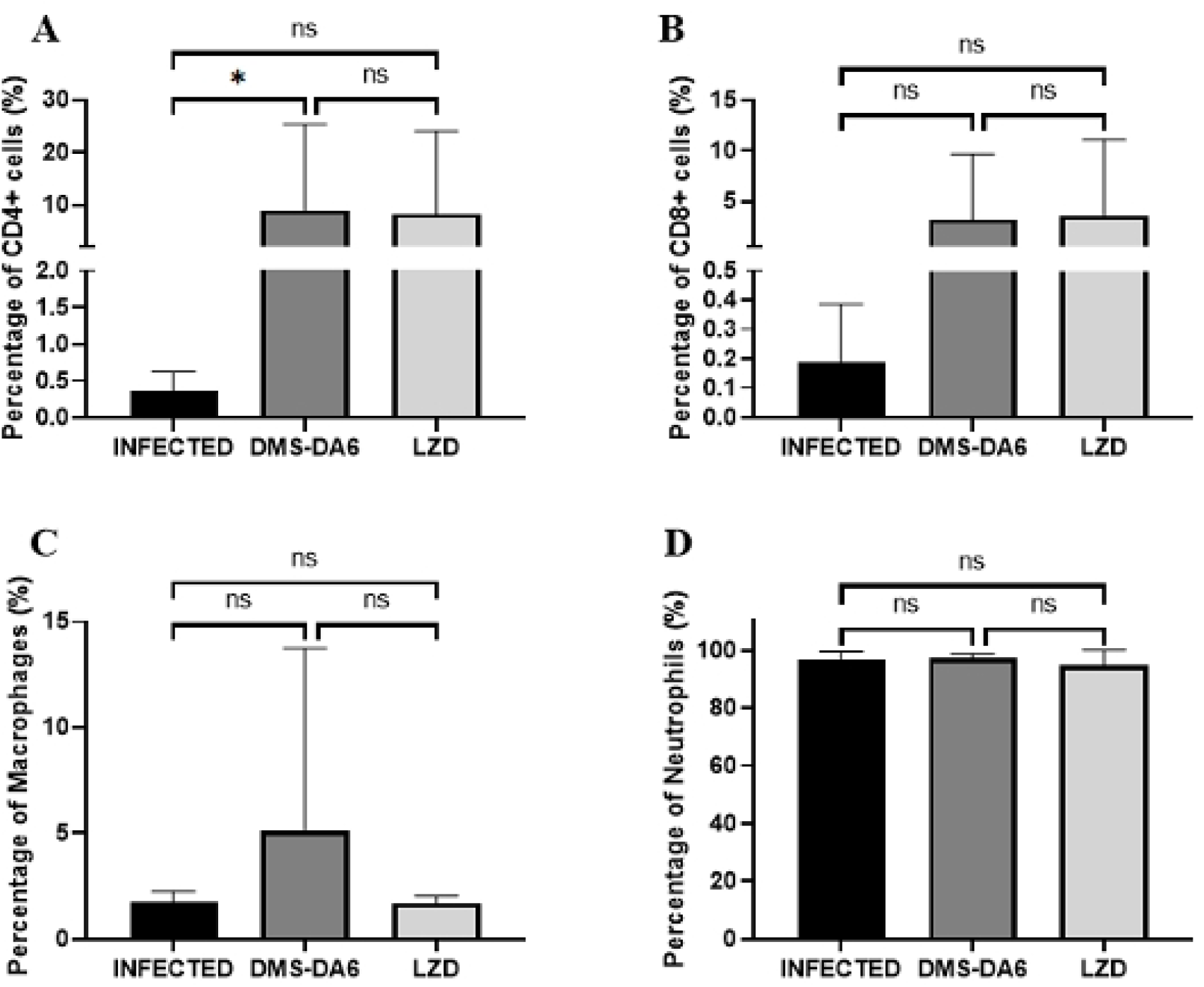
DMS-DA6 treatment boosts the percentage of CD4+ and CD8+ T lymphocytes in the infected footpad but does not affect the percentage of macrophages and neutrophils. The proportions of CD4+ T lymphocytes (A), CD8+ T lymphocytes (B), macrophages (C) and neutrophils (D) in the infected footpad treated or not with DMS-DA6 or Linezolid were evaluated by flow cytometry and analyzed with DIVA software. (n=5) Mann-Whitney test and Unpaired t test *p<0.05 **p<0.01.

## Discussion

Currently, host defense peptides are among the most extensively studied molecules, with intense clinical and non-clinical research evaluating them as potential next- generation antibiotics. Many of these HDPs show antimicrobial activity, as well as the ability to modulate immune system responses *in vitro*.

In this study, we demonstrated that dermaseptin DMS-DA6 is highly effective at controlling chronic infection caused by *N. brasiliensis in vivo*, surpassing the antibiotic linezolid. Indeed, a few injections of low doses of the peptide for short periods of time during the chronic stage of infection resulted in a reduction in the inflammation of the footpad, similar to that achieved with daily injections (every twelve hours), high doses, and prolonged periods of administration of linezolid. Interestingly, although DMS-DA6 exhibited poor antibacterial activity *in vitro*, with a high MIC (500 µM) and MBC (1 mM), *in vivo*, we show it can reduce the bacterial load to a greater extent than the conventional antibiotic. This finding suggests that the bactericidal mechanism of action of DMS-DA6 may be more effective *in vivo* than that of the antibiotic. While linezolid exerts its antibacterial effect by impeding bacterial protein synthesis through binding to ribosomal RNA, DMS-DA6 interacts rapidly with peptidoglycan and membrane lipids to induce membrane disruption and subsequent bacterial death (13,17).

Furthermore, the direct bactericidal activity of DMS-DA6 could be enhanced by the immunomodulatory properties of the peptide. (Indeed, many peptides isolated from frog species have the capacity to regulate inflammatory processes, either by inducing or inhibiting the secretion of cytokines or by modulating the recruitment of immune system cells (18–20). Here, we show that the treatment with DMS-DA6 decreased the production of IL-10 and increased that of IL-1α and IL-6. IL-10 has been reported to be essential in the pathogenic mechanism of intracellular pathogens (15). In *N. brasiliensis* or *Mycobacterium tuberculosis* infections, the interaction of the pathogen with macrophages or dendritic cells induces the differentiation of these cells into foamy cells, which in turn increases the production of IL-10 and the persistence of the bacteria (4). In leishmania infections, the blockade of IL-10 is associated with a decrease in the bacterial load and reduced lesion damage (21,22). At the chronic stage of infections by *Nocardia serolae*, which occur in fish, apoptosis and the phagocytic functions of foamy macrophages have been shown to be inhibited (23). Since IL-10 inhibits activation, apoptosis, and phagocytosis in macrophages, we could hypothesize that by decreasing the production of IL-10 during the chronic stage of infection with *Nocardia brasiliensis*, DMS-DA6 could reactivate infected foamy macrophages and restore apoptosis to potentiate the elimination of the bacteria (15). This is consistent with the observation that DMS-DA6 also increased the levels of IL-1 and IL-6, two cytokines implicated in the activation of Nocardia-infected macrophages.

The increase in the abundance of CD4+ and CD8+ T lymphocytes at the lesion site, resulting from treatment with DMS-DA6, also correlates with the effect observed on cytokines. Indeed, it has been reported that the blockade of the IL-10 axis increased the percentage of CD4+ and CD8+ T lymphocytes in infections induced by *Mycobacterium tuberculosis* (15). These cells are important in the clearance of the pathogen, thereby contributing to the eradication of the infection.

This study is the first to show that treatment with a dermaseptin modulates the leukocyte populations and cytokine production in infected tissue of a living organism and that this modulation may increase the direct bactericidal activity. While the decrease in inflammation in the footpad at the chronic stage of infection can likely be attributed to the immunomodulatory effects of the DMS-DA6 peptide, the greater reduction in bacterial load compared to that observed with linezolid is possibly due to the combination of the direct antibacterial effect and the immunomodulatory capacity of DMS-DA6. In contrast to the available research indicating that linezolid modulates the immune response (24,25), our observations with this infection model did not show such effects. This discrepancy may be attributed to the fact that most of the research has been conducted on acute infections and not on chronic infections, and that such effects have been primarily evaluated over brief periods.

Due to their peptidic nature and bactericidal mechanisms, HDPs offer certain advantages over conventional antibiotics. Specifically, it has been demonstrated that because of their direct membrane interaction and permeabilization, HDPs are less likely to induce bacterial resistance (10). Furthermore, as they are peptidic by nature, they can be degraded by endopeptidases present in the circulation, thus limiting their dispersion in non-targeted tissue and reducing the possibility for severe adverse effects by minimizing their accumulation in these non-target tissues. Collectively, these findings suggest that the antimicrobial peptide DMS-DA6 has the potential to function as an effective therapeutic option for managing actinomycetoma infection caused by *N. brasiliensis*.

## Bibliography

1. Bonifaz A, Flores P, Saúl A, Carrasco-Gerard E, Ponce RM. Treatment of actinomycetoma due to Nocardia spp. with amoxicillin? clavulanate. Br J Dermatol. 2007 Feb;156(2):308–11.

2. Welsh O, Salinas-Carmona MC, la Garza JACD, Rodriguez-Escamilla IM, Sanchez-Meza E. Current Treatment of Mycetoma. Curr Treat Options Infect Dis. 2018 Sep;10(3):389–96.

3. Rosas-Taraco AG, Perez-Liñan AR, Bocanegra-Ibarias P, Perez-Rivera LI, Salinas-Carmona MC. Nocardia brasiliensis Induces an Immunosuppressive Microenvironment That Favors Chronic Infection in BALB/c Mice. Flynn JL, editor. Infect Immun. 2012 Jul;80(7):2493–9.

4. Meester I, Rosas-Taraco AG, Salinas-Carmona MC. Nocardia brasiliensis Induces Formation of Foamy Macrophages and Dendritic Cells In Vitro and In Vivo. Coers J, editor. PLoS ONE. 2014 Jun 17;9(6):e100064.

5. Welsh O, Vera-Cabrera L, Welsh E, Salinas MC. Actinomycetoma and advances in its treatment. Clin Dermatol. 2012 Jul;30(4):372–81.

6. Davidson N, Grigg MJ, Mcguinness SL, Baird RJ, Anstey NM. Safety and Outcomes of Linezolid Use for Nocardiosis. Open Forum Infect Dis. 2020 Apr 1;7(4):ofaa090.

7. Moylett EH, Pacheco SE, Brown-Elliott BA, Perry TR, Buescher ES, Birmingham MC, et al. Clinical Experience with Linezolid for the Treatment of Nocardia Infection. Clin Infect Dis. 2003 Feb;36(3):313–8.

8. Fahal AH. Management of mycetoma. Expert Rev Dermatol. 2010 Feb;5(1):87–93.

9. Drayton M, Deisinger JP, Ludwig KC, Raheem N, Müller A, Schneider T, et al. Host Defense Peptides: Dual Antimicrobial and Immunomodulatory Action. Int J Mol Sci. 2021 Oct 16;22(20):11172.

10. Spohn R, Daruka L, Lázár V, Martins A, Vidovics F, Grézal G, et al. Integrated evolutionary analysis reveals antimicrobial peptides with limited resistance. Nat Commun. 2019 Oct 4;10(1):4538.

11. Pantic J, Jovanovic I, Radosavljevic G, Arsenijevic N, Conlon J, Lukic M. The Potential of Frog Skin-Derived Peptides for Development into Therapeutically-Valuable Immunomodulatory Agents. Molecules. 2017 Dec 13;22(12):2071.

12. Bartels EJH, Dekker D, Amiche M. Dermaseptins, Multifunctional Antimicrobial Peptides: A Review of Their Pharmacology, Effectivity, Mechanism of Action, and Possible Future Directions. Front Pharmacol. 2019 Nov 26;10:1421.

13. Cardon S, Sachon E, Carlier L, Drujon T, Walrant A, Alemán-Navarro E, et al. Peptidoglycan potentiates the membrane disrupting effect of the carboxyamidated form of DMS-DA6, a Gram-positive selective antimicrobial peptide isolated from Pachymedusa dacnicolor skin. Mukherjee A, editor. PLOS ONE. 2018 Oct 16;13(10):e0205727.

14. Salinas-Carmona MC, Torres-Lopez E, Ramos AI, Licon-Trillo A, Gonzalez-Spencer D. Immune Response to Nocardia brasiliensis Antigens in an Experimental Model of Actinomycetoma in BALB/c Mice. Infect Immun. 1999 May;67(5):2428–32.

15. Steffy K, Ahmed A, Srivastava S, Mukhopadhyay S. An Insight into the Role of IL-10 and Foamy Macrophages in Infectious Diseases. J Immunol. 2024 Dec 15;213(12):1729–37.

16. Hancock REW, Sahl HG. Antimicrobial and host-defense peptides as new anti-infective therapeutic strategies. Nat Biotechnol. 2006 Dec;24(12):1551–7.

17. Hashemian SM, Farhadi T, Ganjparvar M. Linezolid: a review of its properties, function, and use in critical care. Drug Des Devel Ther. 2018 Jun;Volume 12:1759–67.

18. Pantic JM, Radosavljevic GD, Jovanovic IP, Arsenijevic NN, Conlon JM, Lukic ML. In vivo administration of the frog skin peptide frenatin 2.1S induces immunostimulatory phenotypes of mouse mononuclear cells. Peptides. 2015 Sep;71:269–75.

19. He Y, Ruan S, Liang G, Hao J, Zhou X, Li Z, et al. A Nonbactericidal Anionic Antimicrobial Peptide Provides Prophylactic and Therapeutic Efficacies against Bacterial Infections in Mice by Immunomodulatory–Antithrombotic Duality. J Med Chem. 2024 May 9;67(9):7487–503.

20. He Y, Shen Y, Feng X, Ruan S, Zhao Y, Mu L, et al. Tree Frog-Derived Cathelicidin Protects Mice against Bacterial Infection through Its Antimicrobial and Anti-Inflammatory Activities and Regulatory Effect on Phagocytes. ACS Infect Dis. 2023 Nov 10;9(11):2252–68.

21. Anderson CF, Lira R, Kamhawi S, Belkaid Y, Wynn TA, Sacks D. IL-10 and TGF-β Control the Establishment of Persistent and Transmissible Infections Produced by Leishmania tropica in C57BL/6 Mice. J Immunol. 2008 Mar 15;180(6):4090–7.

22. Murray HW, Flanders KC, Donaldson DD, Sypek JP, Gotwals PJ, Liu J, et al. Antagonizing Deactivating Cytokines To Enhance Host Defense and Chemotherapy in Experimental Visceral Leishmaniasis. Infect Immun. 2005 Jul;73(7):3903–11.

23. Liu W, Deng Y, Tan A, Zhao F, Chang O, Wang F, et al. Intracellular behavior of Nocardia seriolae and its apoptotic effect on RAW264.7 macrophages. Front Cell Infect Microbiol. 2023 Feb 28;13:1138422.

24. Sauer A, Peukert K, Putensen C, Bode C. Antibiotics as immunomodulators: a potential pharmacologic approach for ARDS treatment. Eur Respir Rev. 2021 Dec 31;30(162):210093.

25. Wang J, Xia L, Wang R, Cai Y. Linezolid and Its Immunomodulatory Effect: In Vitro and In Vivo Evidence. Front Pharmacol. 2019 Nov 28;10:1389.

